# S RNA Intergenic Deletions Drive Viral Interference during Arenavirus Infections

**DOI:** 10.1101/2023.10.31.564889

**Authors:** Matthew Hackbart, Carolina B. López

## Abstract

Arenaviruses, a family of negative-sense RNA viruses spread by rodents, are a leading cause of severe hemorrhagic fever in humans. Due to a paucity of antivirals and vaccines for arenaviruses, there is a need to identify new mechanisms for interfering with arenavirus replication. In several negative-sense RNA viruses, natural viral interference results from the production of non-standard viral genomes (nsVGs) that activate the innate immune system and/or compete for essential viral products. Although it is well established that arenaviruses produce strong interfering activities, it is unknown if they produce interfering nsVGs. Here we show that arenaviruses produce deletions within the intergenic region of their Small (S) RNA genome, which prevents the production of viral mRNA and protein. These deletions are more abundant when arenaviruses are grown in high-interfering conditions and are associated with inhibited viral replication. Overall, we found that arenaviruses produce internal deletions within the S RNA intergenic region that are produced by arenaviruses and can block viral replication. These natural arenavirus interfering molecules provide a new target for the generation of antivirals as well as an alternative strategy for producing attenuated arenaviruses for vaccines.

**AUTHOR SUMMARY:** Arenaviruses are hemorrhagic fever-causing pathogens that infect millions of people a year. There are currently no approved antivirals that target arenaviruses and understanding natural mechanisms that inhibit arenavirus replication is crucial for the development of effective therapeutics. Here, we identify multiple deletions within arenavirus genomes that are associated with the inhibition of viral replication. We show that these deletions prevent viral protein production through the removal of the intergenic region of the viral genome. These deletions were found in all arenaviruses tested in this study representing a novel mechanism for development of new antivirals and vaccines that broadly target the arenavirus family.

## INTRODUCTION

Arenaviruses are negative-stranded, bi-segmented RNA viruses that are a major causative agent of hemorrhagic fever globally. For example, Lassa Virus causes 2 million cases and 5,000-10,000 deaths annually (1, 2, 3, 4). Although arenaviruses are endemic in Africa and South America, there is only one approved vaccine which targets Junin Virus (JUNV) and no targeted antivirals (5). Discovering new mechanisms for attenuating arenavirus infections is imperative for the development of antivirals and vaccines.

One natural mechanism for inhibiting viral replication occurs during viral interference. Viral interference occurs when a virus inhibits its own replication or another virus replication through induction of innate immune responses, such as interferons, modulation of cellular receptors, inhibition of viral protein translation, or by competing for essential viral proteins (6, 7, 8, 9). Viral interference during negative-sense RNA virus infections is associated with the production of non-standard viral genomes (nsVGs), also known as defective viral genomes (DVGs). NsVGs can be deletion viral genomes (delVGs) or copy-back viral genomes (cbVGs) depending on the nature of the genomic rearrangement (Fig 1A). For example, paramyxoviruses will produce cbVGs that can compete for viral polymerases, activate host innate immune sensors, and drive persistent infections (7, 8).

**Figure 1:**
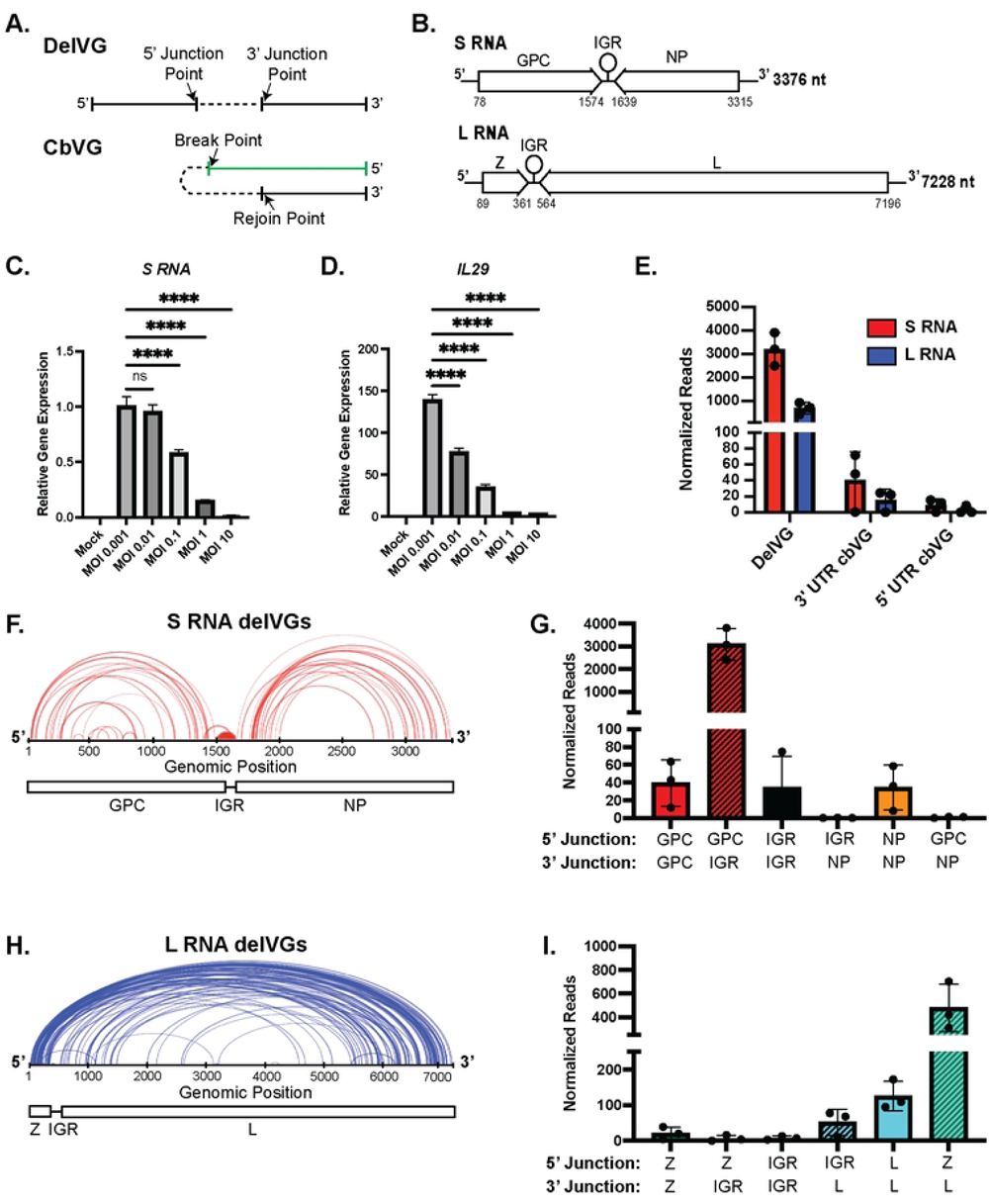
The most consistent nsVGs produced during LCMV infection are S RNA delVGs. A) Schematic of a deletion viral genome (delVG) with the junction points indicated by arrows and copy-back viral genome (cbVG) with the break and rejoin point indicated by arrows. B) Schematic of the LCMV genomes. Z: Matrix Protein; IGR: Intergenic Region; L: Polymerase; GPC: Glycoprotein Complex; NP: Nucleoprotein. C) qPCR of the viral S RNA in A549 Cells were infected with LCMV Armstrong at indicated MOIs. D) qPCR of IL-29 in A549 Cells were infected with LCMV Armstrong at indicated MOIs. qPCR data are a representative of 3 independent repeats with similar results. One-way ANOVA was performed for statistical analysis. ns: p>0.05; ****:p<0.0001. E-J) A549 cells were infected with LCMV Armstrong at an MOI of 0.1. RNA was collected at 48 hpi, sequenced, and analyzed by the VODKA2 pipeline. Reads are normalized as nsVG reads per 1 million viral reads. E) Quantification of nsVG species for the S RNA (red) and L RNA (blue). F) Representation of 5’ junction and 3’ junction points of individual S RNA delVGs. Each arc represents an individual delVG species with the width of the arc proportional to the normalized reads detected per delVG species. Data are a representative of 3 independent sequencing experiments with similar results. G) Quantification of S RNA delVGs that contain 5’ and 3’ junctions in indicated genomic regions. Data are 3 independent sequencing experiments. H) Representation of 5’ junction and 3’ junction points of individual L RNA delVGs. Each arc represents an individual delVG species with the width of the arc proportional to the normalized reads detected per delVG species. Data are a representative of 3 independent sequencing experiments with similar results. I) Quantification of L RNA delVGs that contain 5’ and 3’ junctions in indicated genomic regions. Data are 3 independent sequencing experiments.

Arenaviruses produce two genomic RNA species: the large (L) RNA and small (S) RNA. In ambi-sense fashion, the L RNA encodes a matrix (Z) protein and polymerase (L) protein, while the S RNA encodes the glycoprotein complex (GPC) and a nucleoprotein (NP) (Fig 1B)(10, 11, 12). The S RNA and L RNA each contain an intergenic region (IGR) in between the genes coding for viral proteins. Viral mRNA is produced from the genomic RNA and contains each individual gene and the IGR. The IGRs contain RNA stem loops that function as transcription termination sites for the generation of the viral mRNA and are involved in the viral assembly and budding (13). Arenaviruses that contain genomes with switched IGRs (*e.g.* L RNA containing the S RNA IGR) are severely attenuated *in vivo* (14, 15, 16, 17, 18). In addition, arenaviruses can produce widespread recombination events, such as insertion of multiple IGRs or duplication of gene segments (19), but it is unknown if any of these recombination events can drive viral interference.

Studies with the arenavirus Lymphocytic Choriomeningitis Virus (LCMV) have provided evidence for the presence of nsVGs and viral interference during infection: i) LCMV can produce high levels of defective interfering particles, which block viral replication and spread of LCMV infection (10, 20, 21, 22, 23). ii) Unique RNA species that do not match the viral genomes or mRNA are detectable by northern blot during persistent LCMV infection (24). iii) LCMV infections can produce viral RNA containing small deletions of the genomic ends (25), which may be products of the cap-snatching activity of the viral polymerase (26). iv) Arenavirus vaccine candidates produce differential viral RNA species that contain large internal deletions (14).

With the goal of further understanding arenavirus interference, we sought to identify nsVG species produced naturally during arenavirus infection and determine if and how these nsVGs interfere with virus replication. We show that LCMV produces internal deletion viral genomes (delVGs) in the S RNA IGR that cause viral interference. Utilizing RNA-sequencing and our bioinformatics pipeline, VODKA2, we found that LCMV, JUNV, and Parana Virus (PARV) produce cbVGs and delVGs in both the S and L RNAs. We observed that the most abundant delVGs cover the S RNA IGR and then detected these specific delVGs with RNA Fluorescent *in situ* hybridization. To determine which of the nsVGs are associated with viral interference, we generated two viral stocks: one virus passaged at the standard growth condition and one virus passaged at high multiplicity of infection (MOI) to enhance interfering RNAs. We found that S RNA IGR-delVGs are increased in abundance in the viral stock with viral interference, correlating IGR-delVGs with inhibition of viral replication. Using a minigenome system to determine the mechanism of viral interference, we found that the IGR-delVGs decrease the production of viral mRNA and viral protein. Overall, we found novel delVGs in the LCMV S RNA IGR that can inhibit viral replication through inhibition of viral mRNA and protein production. These IGR-delVGs can be harnessed for antiviral development as well as for the development of attenuated viruses to be used in vaccines.

## RESULTS

### LCMV predominantly produces delVGs

To determine if naturally produced nsVGs can inhibit arenavirus replication, we first characterized the types and quantities of nsVGs that are produced during arenavirus infections. To start, we infected BHK21 cells at an MOI of 0.001 for 72 hours to prepare a stock of LCMV Armstrong 53b (LCMV-Arm) that was capable of replicating to high viral titers according to published protocols (27). To test if interference occurred, we infected A549 lung epithelial cells with LCMV-Arm at increasing MOIs. We observed a decrease in viral RNA at 48 hours post infection (hpi) as the MOI increased, despite no indication of cell death (Fig 1C and Fig S1A+B). Differently from interference observed in other negative-sense RNA viruses, including paramyxoviruses and pneumoviruses, reduction of LCMV RNA did not associate with an increased antiviral response (Fig 1D).

As interference in RNA viruses is generally associated with the presence of nsVGs, we next looked for production of nsVG species at 48 hpi for an MOI of 0.1, when the virus produces high levels of viral RNA but shows the phenotype of viral interference. We used next-generation sequencing to identify nsVGs using our custom bioinformatics pipeline, VODKA2, from LCMV-infected cells (28). We defined specific species of delVGs by the position of the 5’ nucleotide of the junction and the 3’ nucleotide of the junction (Fig 1A). Since LCMV is an ambi-sense virus, we also analyzed cbVGs that contained either the 3’ UTR or 5’ UTR of the viral genome. The cbVGs species are defined by the theoretical break and rejoin points of the recombination event as previously described (Fig 1A)(7, 8). We found that LCMV-Arm produces cbVGs containing the 3’ UTR or 5’ UTR, and a majority of the nsVGs found were delVGs (Fig 1E). Both the S RNA and L RNA genomic segments produced delVGs with distinct distributions of junction points (Fig 1F+H). The S RNA had 1 major set of abundant delVGs that removes the end of the GPC gene and part of the IGR. The S RNA also contains minor deletions that removes either portions of the GPC gene, the IGR, or the NP gene. (Fig 1F+G). Most of the L RNA delVG species contain 5’ junction points occurring in the first 500 nucleotides within the Z gene and 3’ junction points occurring after 5000 nucleotides of the genome, thereby deleting most of the L polymerase gene (Fig 1H+I). Minor L RNA delVGs also occur within the L gene and the IGR-L segment. Interestingly, both S RNA and L RNA delVGs tend to remove the IGRs of the genomic RNAs, which are necessary for viral replication (13). Unlike the delVGs, the cbVG species are more evenly distributed throughout the genome (Fig S2A+B).

The most abundant delVG species detected by RNA-Seq are deletions in the S RNA that contain 5’ junction points within the end of the GPC gene and 3’ junction points within the IGR stem-loop structure (Table S1 and Fig 2A). While the 5’ junction points vary, the most consistent 3’ junction point is around nucleotide 1613 in the IGR. The most abundant of these S RNA IGRs are IGR-delVG 1560_1613 and IGR-delVG 1572_1613, each having about 1500 reads per 1 million viral reads detected (Fig 2B). To test if the IGR-delVGs were specific to LCMV-Arm, we also sequenced a stock of LCMV-Clone 13 (LCMV-Cl13) and found the dominant delVG species were also delVG 1560_1613 and delVG 1572_1613 (Fig S3A+B). The IGR-delVGs can also be detected by RT-PCR over the LCMV-Arm S RNA IGR (Fig S4A and Table S2).

**Figure 2:**
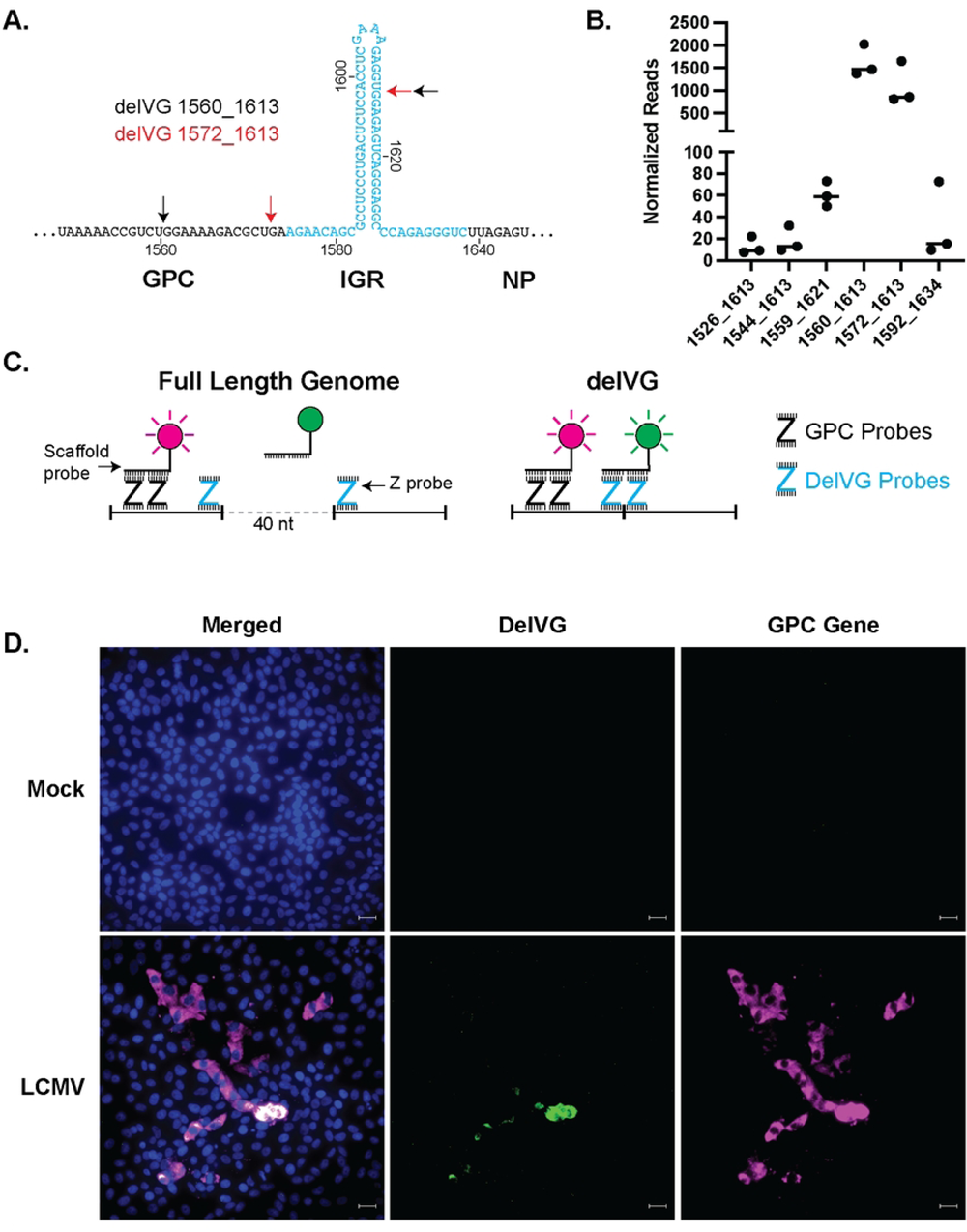
The most abundant S RNA delVGs delete the GPC-IGR. A) Schematic for location of the conserved S RNA IGR-delVGs. The IGR is highlighted in blue. Black arrows indicate the junction points for delVG 1560_1613 and red arrows indicate the junction points for delVG 1572_1613. B) Quantification of the individual IGR-delVGs. Data are 3 independent sequencing experiments. C) Schematic for RNAscope with Z probes binding to the wildtype or delVG genomes. GPC Gene probes produce a fluorescent signal (magenta) upon binding to the wildtype and delVG RNAs, whereas the delVG probes only produce a fluorescent signal (green) when binding to the delVG RNA. D) Vero E6 cells were infected with LCMV Armstrong at an MOI of 0.1. Images were processed by RNAscope and stained for nuclei (blue), delVG 1572_1613 (green) and GPC gene (magenta). Scale bar is 20 uM. Images are representative of 3 independent imaging experiments.

### Major S RNA delVGs can be detected in infected cells using a PCR-independent strategy

As the IGRs play critical roles during viral replication, the abundance of IGR deletions prompted us to hypothesize that IGR-delVGs generated during virus replication are involved in LCMV interference. However, during validation of our RNA-seq analysis pipeline, we learned that the IGR-delVGs can be detected from an *in vitro-*transcribed RNA that contained the GPC gene, IGR, and an mCherry gene replacing the NP gene (Fig S4B+C). Detection of IGR-delVGs in the absence of viral replication proteins suggests that IGR-delVGs can be generated by either the T7 RNA polymerase, RT enzyme, or subsequent PCRs in the RNA-seq library preparation and we needed to validate their presence during infection using T7 and RT-PCR independent methods.

To do this, we took advantage of Basecope Technology (ACD Biosystems), an RNA fluorescent *in situ* hybridization (FISH) assay that utilizes paired DNA “Z”-probes and amplifying scaffolding probes to produce a detectable fluorescent signal. Specifically, we used probes to detect the S RNA GPC Gene and the delVG 1572_1613. The GPC probes, which serve as a marker for viral RNA, bind to the genome allowing the scaffolding probe to bind a horse radish peroxidase and a fluorescent signal to be amplified with tyramide signal amplification (Fig 2C). In the full-length standard viral genomes, the delVG Z-probes are separated by 40 nucleotides so the scaffolding probes cannot efficiently bind to the two delVG Z-probes. However when delVG 1572_1613 is generated, the 40 nucleotides are removed and the Z-probes are able to bind and hybridize with the scaffolding probe, producing a fluorescent signal (Fig 2C) To test this assay, we transfected plasmids that transcribe either the wildtype GPC gene or the delVG RNA. While the RNA-FISH probe against the GPC gene is detected for both transfections, the delVG probe set only produces a fluorescent signal when the delVG RNA is present (Fig S5) During viral infection, the delVG RNA was detectable in a subset of infected cells, confirming that delVG 1572_1613 is produced during LCMV infection (Fig 2D).

### High levels of S RNA IGR-delVGs are found in high-interfering infections

We next tested if IGR-delVGs are enhanced in infections with LCMV stocks that have high-interfering activity. When comparing infections with increased MOIs, we observed an increase in delVG 1572_1613 at higher MOIs (Fig S1C). To compare viral interference during infections at the same MOI, we generated different stocks of LCMV grown by either low MOI passaging (LCMV-LMP) or high MOI passaging (LCMV-HMP)(Fig 3A). As expected, subsequent infections in BHK21 cells showed decreased production of infectious particles during LCMV-HMP infection compared to LCMV-LMP (Fig 3B+C). This interference did not associate with an enhanced early antiviral interferon response during infection of A549 cells (Fig S6A-C). Upon next generation sequencing and VODKA2 analysis of the passaged stocks, we discovered that the LCMV-HMP stock had an increased abundance of the IGR-delVGs, but no increase in other nsVGs (Fig S7A+B). To determine if the increase in delVGs were not a product of the higher MOI for the LCMV-HMP stock, we determined if LCMV-HMP had increased IGR-DelVGs during subsequent infections at the same MOI. Upon infection of Vero E6 cells with either the LCMV-LMP or LCMV-HMP, we again observed viral interference of the production of infectious particles during LCMV-HMP infection (Fig 3D). Importantly, LCMV-HMP had increased abundance of IGR-delVG-positive cells at 24 hpi (Fig 3E+F), while the LCMV-LMP infection eventually produced IGR-delVG-positive cells detectable at 48 hpi. We also observed an increase in IGR-delVGs at 24 hpi during High MOI infection by qPCR (Fig S6D). These data indicate that LCMV with increased IGR-delVGs exhibits higher levels of viral interference without enhancing activation of the interferon pathway.

**Figure 3:**
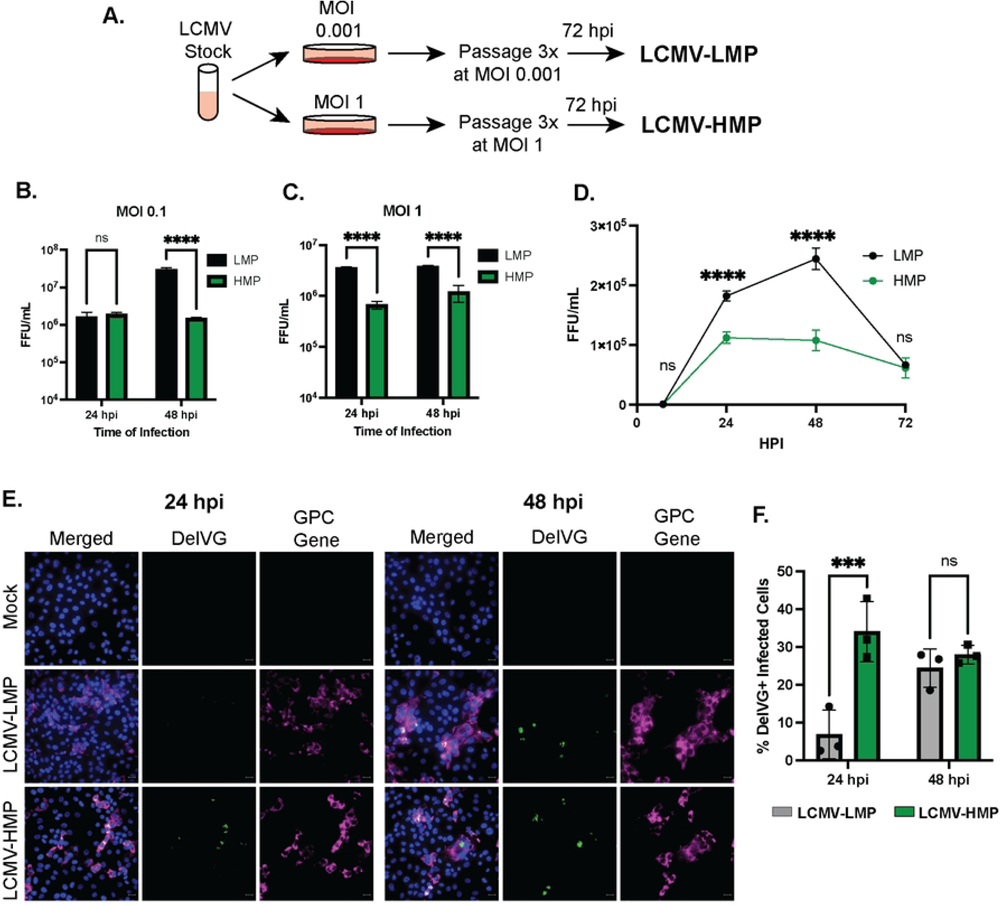
S RNA IGR-delVGs are increased during LCMV interference. A) Schematic for the generation of LCMV stocks grown at the standard MOI of 0.001 (LCMV-LMP) and at high MOI of 1 (LCMV-HMP). B+C) BHK21 cells were infected with either LCMV-LMP or LCMV-HMP at an MOI of B) 0.1 or C) 1. Cellular supernatant was collected at indicated timepoints and focus-forming units (FFU) were quantified. Data are 3 biological replicate infections. Two-way ANOVA was performed for statistical analysis. ns: p>0.05; ****: p<0.0001. D) Vero cells were infected with either LCMV-LMP or LCMV-HMP at an MOI of 1. Cellular supernatant was collected at indicated timepoints and focus-forming units (FFU) were quantified. Data are 3 biological replicate infections. Two-way ANOVA was performed for statistical analysis. ns: p>0.05; ****: p<0.0001. E) Vero cells were infected with either LCMV-LMP or LCMV-HMP at an MOI of 1. Cells were processed at 24 hpi or 48 hpi with RNAscope for detection of nuclei (blue), delVG 1572_1613 (green), and GPC gene (magenta). Scale bar is 20 uM. Images are representative of 3 independent imaging experiments. F) Quantification of the percentage of LCMV-infected cells that were delVG-positive. 2-way ANOVA was performed for statistical analysis. ns: p>0.05; **: p<0.01. Data are 3 independent imaging experiments.

### The S RNA IGR-delVGs reduce protein production during minigenome replication

The arenavirus IGRs were previously shown to function as transcription termination sequences during the production of viral mRNAs. Minigenome replication systems that replicate the S RNA genome by co-expression with the NP and L proteins show that complete deletion of the S RNA IGR led to a loss of viral mRNAs and a decrease in viral protein production (13). We therefore hypothesized that our observed IGR-delVGs similarly inhibit the production of viral mRNA and protein production.

Utilizing an LCMV minigenome system, we transfected a wildtype minigenome or minigenomes containing deletions 1560_1613 or 1572_1613 (Fig 4A). Upon co-transfection with the NP and L proteins, the wildtype minigenome produced a strong mCherry signal while the delVG-containing minigenomes showed markedly reduced mCherry signal, suggesting reduced protein production in the delVG conditions (Fig 4B). The decrease in protein production was not due to decreased replication of the delVG minigenomes. We instead detected increased RNA levels of the genomic RNA and GPC gene when the IGR-delVGs are present (Fig 4C). When quantifying the proportion of viral mRNA to genome, we find that the viral mRNA is decreased during delVG minigenome replication compared to the wildtype (Fig 4D). These data suggest that while the IGR-delVG minigenome may have higher replication rates compared to the wild-type minigenome, the delVG minigenomes have decreased mRNA production relative to genomic RNA levels and decreased total protein production compared to wildtype.

**Figure 4:**
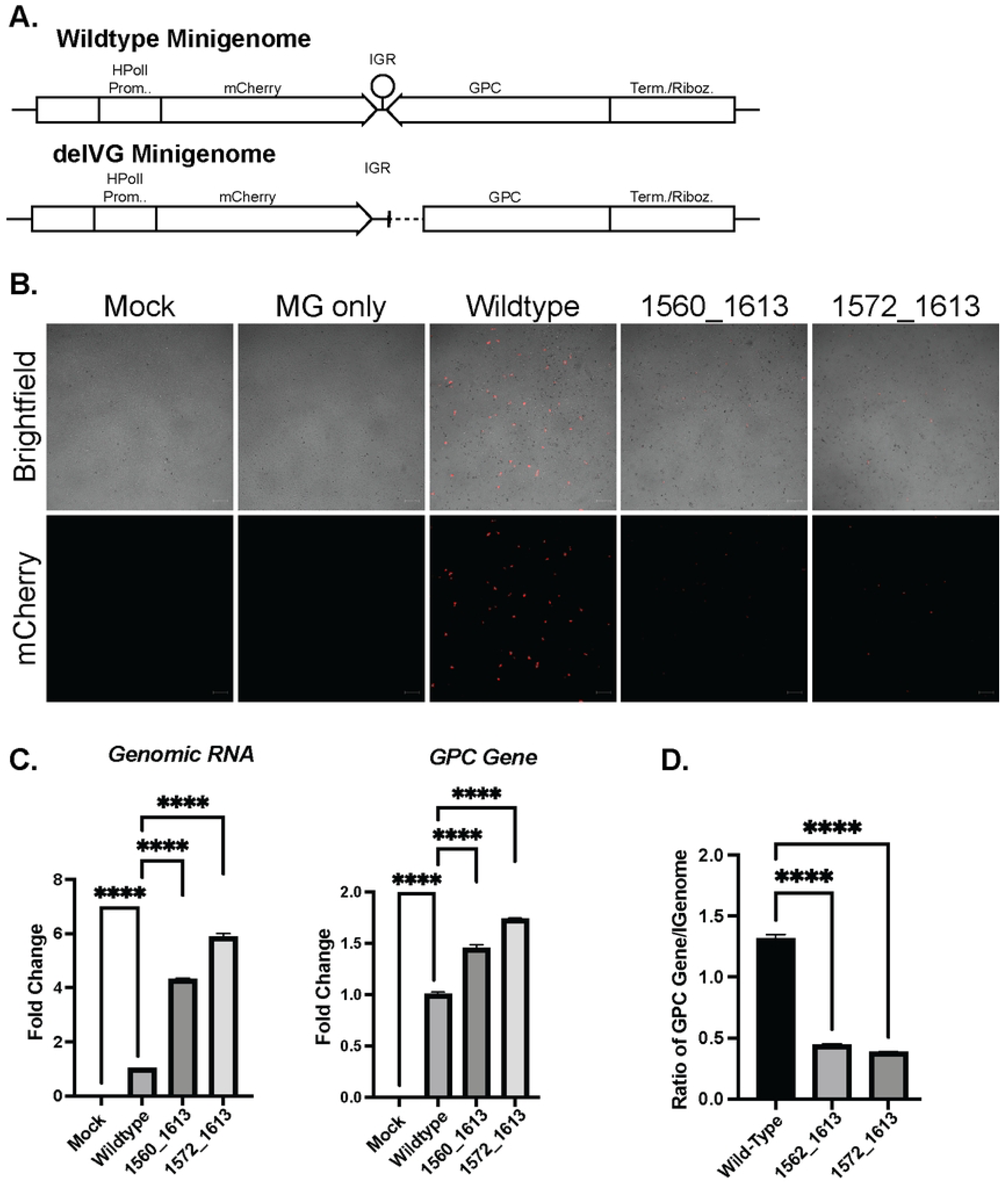
The S RNA IGR-delVGs decrease viral RNA and protein production. A) Schematic of LCMV minigenomes (MG) that contain the human PolI promoter, mCherry gene, intergenic region (IGR), glycoprotein complex, and termination/ribozyme sequence. The delVG minigenomes contain either the 1560_1613 or 1572_1613 deletions. B) Vero E6 cells were co-transfected with the MG plasmid, NP helper plasmid, and L helper plasmid. At 24-hour post transfection, cells were imaged for mCherry signal. Scale bar is 200 uM. Images are representative of 3 independent imaging experiments. C) RNA was collected at 24 hpt from cells transfected with wildtype or delVG MGs and qPCR was performed to measure Genomic RNA and GPC Gene. D) The ratio of GPC Gene to Genomic RNA was calculated. One-way ANOVA was performed for statistical analysis. ****: p<0.0001. Data are representative of 3 independent experiments with similar results.

### JUNV and PARV produce S RNA IGR-delVGs

Lastly, we sought to determine if other arenaviruses produced similar IGR-delVGs as LCMV. We infected Vero E6 cells with Junin Virus Candid 1 (JUNV) or Parana Virus (PARV) for 48 hours. RNA was collected, sequenced, and analyzed for nsVGs using the VODKA2 analysis pipeline. Both JUNV (Fig 5A) and PARV (Fig 5D) produced delVGs in the S RNA and L RNA, while very few or no cbVGs were detected. JUNV produced four S RNA IGR-delVG species with the most abundant being delVG 1554_1582 (Fig 5B+C). PARV also produced four species of S RNA IGR-delVGs with delVG 1607_1649 being most detected (Fig 5E+F). Interestingly, the IGR-delVGs of JUNV and PARV again appear to partially remove the IGR, similar to the LCMV delVGs. The locations of the deletion junctions within the IGR stem loops for PARV, JUNV, and LCMV suggests that there may be a conserved mechanism for the generation of these delVGs in arenaviruses.

**Figure 5:**
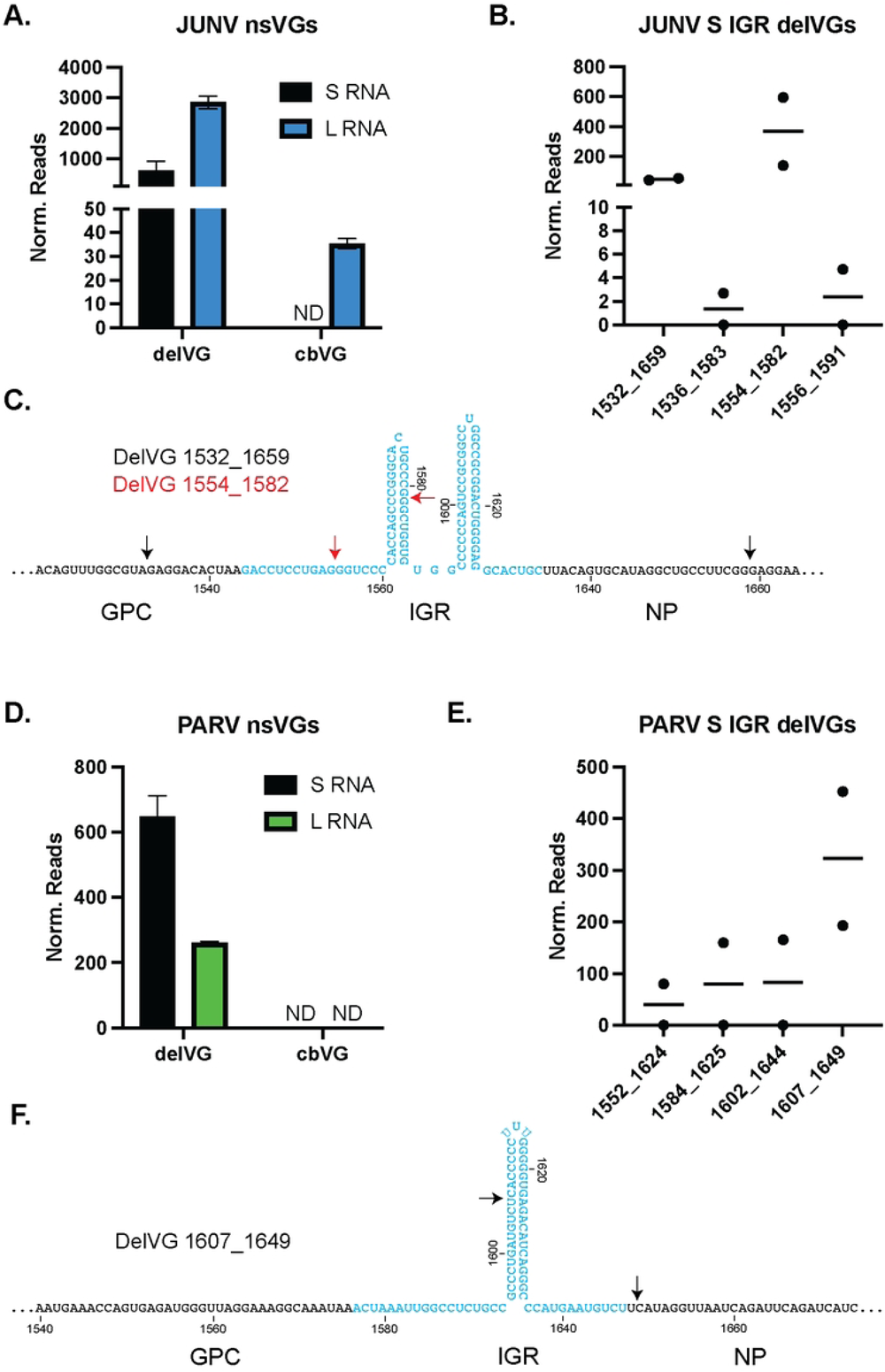
JUNV and PARV infections also produce S RNA IGR-delVGs. Vero E6 cells were infected with either A-C) Junin Virus Candid 1 (JUNV) or D-F) Parana Virus (PARV) at an MOI of 0.1. Cellular RNA was isolated at 48 hpi and RNA-sequencing/VODKA2 analysis was performed. Reads were normalized as nsVG reads per 1 million viral reads. A) Quantification of nsVGs during JUNV infection for the S and L RNAs. B) Quantification of the JUNV S RNA IGR-delVGs C) Schematic representation of the JUNV S RNA IGR-delVGs. Black arrows indicate delVG 1532_1659 and red arrows indicate delVG 1554_1582. D) Quantification of nsVGs during PARV infection for the S and L RNAs. E) Quantification of the PARV S RNA IGR-delVGs F) Schematic representation of the PARV S RNA IGR-delVGs. Black arrows indicate delVG 1607_1649. Data are representative of 2 individual sequencing experiments.

## DISCUSSION

Here, we show that LCMV produces delVGs that can inhibit viral replication. The major species of delVGs are localized to the S RNA IGR and delete out portions of the GPC gene and IGR. Similar delVGs were found in other arenavirus infections, JUNV Candid#1 and PARV. The S RNA IGR-delVGs have increased abundance during higher-interfering LCMV infection, suggesting a mechanism for viral interference. We show that S RNA IGR-delVGs can inhibit the production of viral mRNA and protein, providing a possible mechanism for control of replication.

Important questions remain about how these IGR-delVGs are produced and if the generation of the IGR-delVGs can be modulated. A recent study found that Lassa Virus (LASV) that lacks the nucleoprotein exonuclease activity had increased rates of deletions in both the S RNA and L RNA (29). Interestingly, the major S RNA deletions reported in that study map similarly to the IGR region as we observed for LCMV, JUNV, and PARV. It is not understood how the arenavirus NP exonuclease inhibits the formation of the IGR delVGs, but modulation of this activity through small molecule inhibitors could potentially attenuate arenavirus replication by increasing delVG production. Thus, generation and regulation of arenavirus delVGs represent an important area of further study for development of novel antivirals.

The interference by the IGR-delVGs suggests the possibility of targeting the IGRs for vaccine development. Interestingly, the IGR has already been shown to be an efficient target at attenuating arenaviruses for vaccine generation (14, 15, 16, 17, 18). Using reverse genetics, recombinant arenaviruses that contain the S RNA IGR in both the S RNA and L RNA have been developed. These recombinant arenaviruses are completely attenuated during infection of mice and guinea pigs (17, 18). Since we observe that the S RNA IGR is prone to develop delVGs, it is possible that the L RNA in these recombinant vaccines, which contain the S RNA IGR, have enhanced delVG production, thus preventing protein expression from the L RNA and attenuating the virus. Additionally, other vaccine candidates that show promising attenuation produce truncated viral RNAs (14).

With the rapid development of mRNA vaccines and siRNA-based therapies, utilization of RNA substrates in the clinic is rapidly expanding. IGR-delVGs present an interesting candidate for arenavirus-specific anti-viral therapy, which are currently lacking. We observe that minigenomes containing IGR-delVGs are capable of increased replication compared to the full-length minigenome (Fig 4). It is likely based on these findings, and data from other viruses, that IGR-delVGs delivered to an infected cell will outcompete the normal viral genome from replicating and prevent infectious particle release resulting in control of viral infection and spread. Since we observed similar IGR-delVGs with JUNV and PARV, we would be able to develop similar IGR-delVGs for each arenavirus.

Overall, we have discovered a novel mechanism for attenuation of arenavirus replication. The IGR-delVGs allow the virus to control its own replication by inhibiting the viral protein production. These IGR-delVGs present a novel target for the generation of antivirals and provides a mechanism for the attenuation of many proposed arenavirus vaccines.

## METHODS AND MATERIALS

### Cells and Viruses

BHK-21 cells (hamster kidney cells, ATCC, CCL-10), Vero E6 cells (green monkey kidney cells, ATCC, CRL-1586), and A549 cells (human lung cells, ATCC, CCL-185) were cultured at 37°C and 5% CO2 with Dulbecco’s modified Eagle’s media (Thermofisher, 11995065) supplemented with 10% fetal bovine serum (FBS), 1 mM sodium pyruvate, 2 mM L-Glutamine, and 50 mg/mL gentamicin. All cell lines were treated with mycoplasma removal agent (MP Biomedicals, 093050044) and routinely tested for mycoplasma before use. Lymphocytic choriomeningitis virus (LCMV) Armstrong 53b (Genbank Accession number: L RNA; AY847351.1, S RNA; AY847350.1) and Clone 13 (Genbank Accession number: L RNA; DQ361066.1, S RNA; DQ361065.2) were propagated in BHK-21 cells according to published methods (27). Junin Virus Candid 1 (Genbank Accession number: L RNA; AY819707.2, S RNA; FJ969442.1) and Parana Virus (Genbank Accession number: L RNA; NC_010761.1, S RNA; NC_010756.1) were propagated in Vero E6 cells.

### LCMV infection

Cells were infected with LCMV-Armstrong at indicated MOIs. Briefly, cells were washed with PBS, then infectious media containing virus and no FBS was added for 1 hour with rocking at 37°C and 5% CO2. Infectious media was removed and cell culture media was added. Infected cells were incubated at 37°C and 5% CO2 for indicated timepoints. LCMV was titered by focus forming assay. Briefly, Vero E6 cells were infected with sequential dilutions of LCMV supernatant and overlayed with 0.3% agarose with DMEM containing 5% FBS. At 2 days post infection, cells were fixed with 3.7% formaldehyde, permeabilized with 0.1% Triton X-100, blocked with 5% FBS in PBS, and immunostained for LCMV nucleoprotein. Cells were stained with 1:100 dilution of mouse anti-nucleoprotein (Abcam, AB31774), 1:1,000 dilution of biotinylated goat anti-mouse IgG (Southern Biotech, 1031-08), and 1:1,000 Strep-AP (Invitrogen, 21324). Individual Foci were visualized with NBT/BCIP (Invitrogen, 34042) and quantified.

LCMV-Low MOI Passaged (LMP) and LCMV-High MOI Passaged (HMP) were generated by passaging the LCMV Armstrong stock virus for 3 passages at MOIs of 0.001 and 1, respectively. Viral supernatants were collected at 72 hours post infection for each passage. Viral stocks were sequenced by RNA-sequencing as described below.

### RNA-Sequencing and VODKA2 Analysis

RNA was collected at indicated timepoints and isolated with Trizol (Thermofisher, 15596026) for cell culture or Trizol LS (Thermofisher, 10296010) for viral supernatants according to the manufacturer’s protocols. RNA integrity and concentration were measured by Bioanalyzer (Agilent) and Qubit (Invitrogen) respectively.

Sequencing libraries were prepared with Illumina Truseq Stranded Total RNA kit with Ribozero Gold Depletion (Illumina, 20020599). Libraries were then sequenced for 2 x 150 paired-end reads on either a NovaSeq 6000 (Illumina) with the WUSTL Genome Technology Access Center or Nextseq 550 (Illumina) with the WUSTL Center for Genome Sciences & Systems Biology.

VODKA2 analysis was performed for detection of cbVGs and delVGs as previously published (28). Additional filters were added to exclude background from samples. For cbVGs, species with less than 2 reads and species whose predicted genomes were greater than the wild-type virus genome were excluded from the study. For delVGs, species with less than 2 reads and species with deletions of 5 or less nucleotides removed were excluded from the study.

### RNAscope

Vero E6 cells were either transfected with pCAGGs-GPC-HA or pCAGGs-GPC-Cpep-HA or infected with LCMV-LMP or LCMV-HMP stocks at an MOI of 0.1. At indicated timepoints, cells were fixed with 3.7% formaldehyde, then processed for imaging using the RNAScope Multiplex Fluorescent Detection Kit v2 (ACD Bio, 323110) according to manufacturer’s protocol. GPC Gene probe detects nucleotides 201-243 of S RNA segment. DelVG 1572-1612 probe detects nucleotides 1542-1572 and 1612-1623 of the S RNA. Probes were designed by ACD Bio (acdbio.com). Dyes used for detection were TSA Plus Cy3 System (Akoya Biosciences, NEL744001KT) and TSA Plus Cy5 System (Akoya Biosciences, NEL745001KT). Coverslips were mounted with ProLong Diamond Antifade (Thermofisher, P36961) and imaged using a Zeiss Axio Observer Widefield microscope (Zeiss).

For quantification, approximately 1,000 cells and 200 infected cells were imaged per individual replicate. Images were quantified using Aggrecount automated image analysis (https://aggrecount.github.io) using FIJI software (https://imagej.net/software/fiji/). In brief, cells were counted as infected if they had cell mean fluorescence over the threshold for the GPC probes. Cells were counted as delVG positive if they had cell mean fluorescence over the threshold for the delVG probes.

### RT-PCR and qPCR

RNA was collected at indicated timepoints and isolated with Trizol (Thermofisher, 15596026) for cell culture or Trizol LS (Thermofisher, 10296010) for viral supernatants according to the manufacturer’s protocols. cDNA was generated with random hexamers using the High-Capacity RNA to cDNA kit (Thermofisher, 4387406) according to the manufacturer’s protocols. PCRs were performed with Hot Start Taq DNA Polymerase (NEB, M0495L). IGR delVG bands were separated on a 2% agarose gel. For qPCR quantification, cDNA was quantified with Power SYBR Green Mix (Thermofisher, 4367660). For quantification of delVG 1572_1613, Taqman Universal Master Mix (Thermofisher, 4440040) was used to amplify cDNA generated above. All qPCRs were performed on a Quantstudio 5 PCR system (Thermofisher, A34322).

RT-PCR Probes:

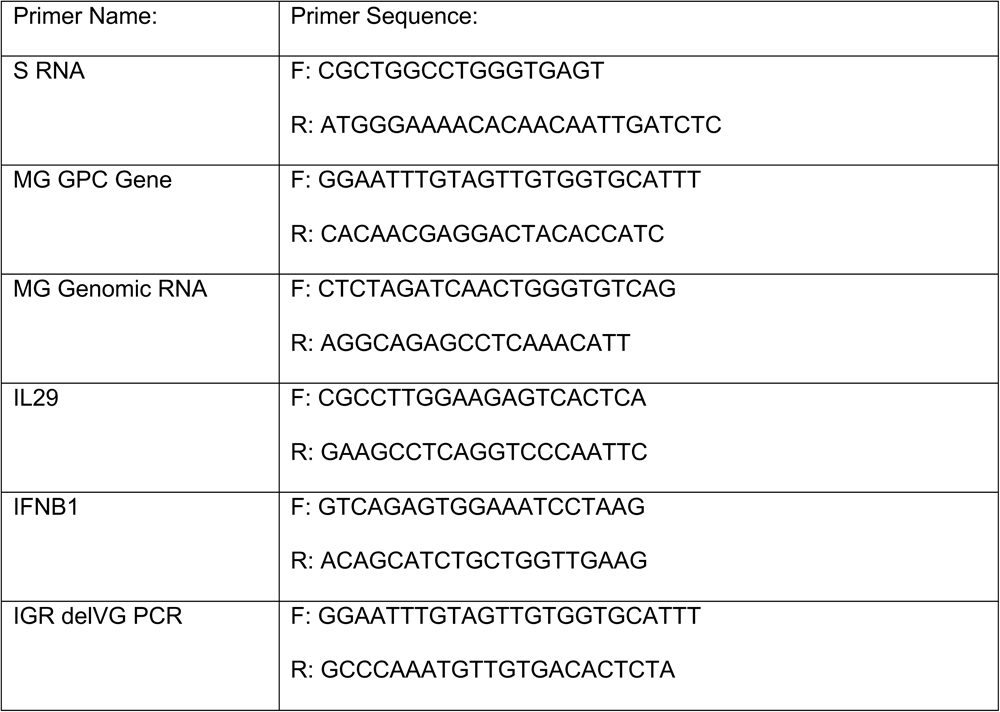

Taqman Primers/Probes:

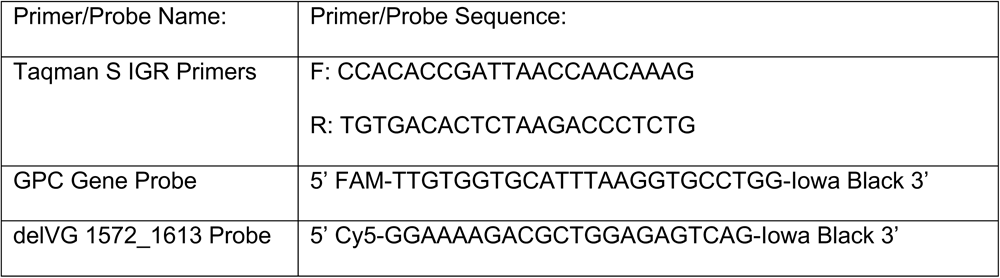

### Minigenome Replicon System

The LCMV S RNA minigenome plasmid and LCMV protein-expression plasmids was kindly gifted by Dr. Luis Martínez-Sobrido (UTMB)(15). The S RNA MG used was cloned to include the full-length GPC and an mCherry gene replacing the NP gene by digestion with AvrII (NEB, R0174S), then insertion gene fragments with NEBuilder HIFI DNA Assembly Mix (NEB, E5520S). For transfection and replication, Vero E6 cells were co-transfected with Minigenome, NP, and L plasmids with Lipofectamine 2000 (Thermofisher, 11668019). Cells were incubated for 24 hours until imaged for mCherry Expression or processed for qPCR.

### In Vitro Transcription of Minigenome RNA

The LCMV S RNA minigenome was cloned into a pSL1180-T7 plasmid. *In vitro* transcription was performed with the Megascript T7 Transcription Kit (Thermofisher, AM1334) and RNA was isolated with LiCl precipitation according to manufacturer’s protocols. Purified RNA was measured by Qubit and tested for quality by Bioanalyzer (Agilent).

### Statistics

All statistics were calculated using GraphPad Prism. Version 9. Specific tests and significance values are indicated in each figure legend.

## ACKNOWLEDGEMENTS

We would like to acknowledge Dr. Luis Martinez-Sobrido for gifting LCMV antibodies and plasmids. We would like to thank Dr. Sean Whelan for gifting stocks of the Junin Virus Candid #1 and Parana Virus. We would also like to acknowledge Emna Achouri for helping with VODKA2 analysis.

## Funding

NIH T32 HL007317-44 (MH)

NIH T32 AI007163-45 (MH)

NIH A137062 (CBL, MH)

WashU BJC Investigator Program (CBL)

## Authors Contributions

Conceptualization: MH, CBL

Methodology: MH

Investigation: MH

Visualization: MH

Supervision: CBL

Writing-Original Draft: MH

Writing-Review and Editing: MH, CBL

Authors declare that they have no competing interests.

All data are available in the main text or the supplementary materials. Raw sequencing files are deposited at NCBI Sequence Read Archive (SRA) with the BioProject ID PRJNA1032960.

